# LM6-M: a high avidity rat monoclonal antibody to pectic α-1,5-L-arabinan

**DOI:** 10.1101/161604

**Authors:** Valérie Cornuault, Fanny Buffetto, Susan E Marcus, Marie-Jeanne Crépeau, Fabienne Guillon, Marie-Christine Ralet, J Paul Knox

## Abstract

1,5-arabinan is an abundant structural feature of side chains of pectic rhamnogalacturonan-I which is a matrix constituent of plant cell walls. The study of arabinan in cells and tissues is driven by putative roles for this polysaccharide in the generation of cell wall and organ mechanical properties. The biological function(s) of arabinan is still uncertain and high quality molecular tools are required to detect its occurrence and monitor its dynamics. Here we report a new rat monoclonal antibody, LM6-M, similar in specificity to the published rat monoclonal antibody LM6 (Willats et al. (1998) Carbohydrate Research 308: 149-152). LM6-M is of the IgM immunoglobulin class and has a higher avidity for α-1-5-L-arabinan than LM6. LM6-M displays high sensitivity in its detection of arabinan in in-vitro assays such as ELISA and epitope detection chromatography and in in-situ analyses.

**Abbreviations:**

Ara: Arabinose
BSA: Bovine Serum Albumin
Gal: Galactose
GalA: Galacturonic acid
EDC: Epitope detection chromatography
ELISA: Enzyme-Linked Immunosorbent Assay
mAb: Monoclonal antibody
PBS: Phosphate-buffered saline
Rha: Rhamnose
RG-I: Rhamnogalacturonan-I

## Introduction

Plant cell walls are highly complex structures composed of load-bearing cellulose microfibrils and interspersed sets of matrix polysaccharides. Pectins, a major component of cell wall matrices, are galacturonic acid-rich polysaccharides and include homogalacturonan and rhamnogalacturonan domains (Caffall and Mohnen, 2009). Pectic rhamnogalacturonan-I (RG-I) is a highly heterogeneous and variable domain of pectin with a rhamnogalacturonan backbone and diverse side chains that are mostly neutral sugars and the major components of which are 1,4-galactosyl and 1,5-arabinosyl residues (Willats et al., 2001; Caffall and Mohnen, 2009). RG-I molecules are known to be structurally highly heterogeneous in cell and developmental contexts and are strongly implicated in influencing the mechanical properties of cell walls and plant materials (Lee et al., 2012; Paniagua et al., 2016; Mikshina et al., 2017) but precise modes of action are not known.

Tracking the fine and subtle changes in RG-I structures in cell and biochemical contexts is a challenging task. In this context monoclonal antibodies (mAbs) are important tools for detection and tracking of polysaccharide structural features or epitopes (usually 3 to 5 sugars) and have great versatility and sensitivity. By nature antibody-based assays amplify signals and are particularly appropriate for the identification of structures at low levels of occurrence (often of pg of material) within the context of complex mixtures of polysaccharides and other molecules. mAbs targeting cell wall polysaccharides are routinely used in immunofluorescence microscopy to identify cell wall locations (Lee et al., 2012; Palmer et al., 2015; Amsbury et al., 2016) and also subsequent to cell wall deconstructions in a range of quantitative and high through-put protocols (Cornuault et al., 2014; Sillo et al., 2016; Baldwin et al, 2017). A widely used mAb that binds to 1,5-L-arabinan motifs in the side chains of RG-I is a rat mAb known as LM6 (Willats et al., 1998). Here we report the isolation (subsequent to immunization with a sugar beet RG-I immunogen) of new rat mAb with a similar specificity to LM6. This probe, LM6-M, is of a different immunoglobulin isotype and has a higher avidity making it a highly effective mAb for pectic arabinan in *in-situ* and *in-vitro* analyses.

## Materials and Methods

### Preparation of sugar beet RG-I oligosaccharides for immunogen preparation

RG-I polysaccharides were extracted from 100 g of sugar beet pulp with 3 L of 0.1 M NaOH at 90 °C for 2 h. The mix was filtered through a 90 μm mesh size nylon cloth and through sintered glass to remove smaller particles and was adjusted to pH 6.5 by addition of 6 M HCl. The extract was concentrated to 1 L by rotary vacuum evaporation at 40°C. Pectins were recovered by precipitation with 2 vol of 97% (v/v) aqueous ethanol overnight at 4°C. The resulting suspension was then centrifuged at 8100 *g* for 15 min. The pellet was recoveredand washed with 97% ethanol. The precipitate was suspended in water and dialyzed against distilled water overnight at 4°C prior to freeze drying. 700 mg of sugar-beet root polysaccharides enriched in pectins were recovered. These were loaded on to a DEAE-Sepharose CL-6B column (27 cm ×26 mm) in three rounds. The column was equilibrated with degassed 50 mM sodium acetate buffer pH 4.5 at a flowrate of 1.5 ml/min. The elution gradient used to separate out the polysaccharides was in two steps: 0 to 450 mL - 50 mM sodium acetate pH 4.5; 450 mL to 1800 mL 0 to 100% 50 mM sodium acetate + 0.6 M NaCl. The fractions were collected and fractions were analysed as described in Cornuault et al., 2015. The fractions containing RG-I were pooled together, dialyzed and freeze-dried. RG-I-enriched polysaccharides were then chemically hydrolysed using 100 mM HCl at 85°C for 1.5 h in order to reduce the extent of side chains and expose a range of epitopes. The mixture was concentrated to a volume of 80 mL by rotary vacuum evaporation and the pH was adjusted to 7 using 100 mM NaOH, prior to dialysis against distilled water and freeze-drying. RG-I backbones were then digested using a rhamnogalacturonase (Ralet et al., 2010) as follows: RG-I preparation (1 mg/mL) was dissolved in 10 mM sodium acetate buffer, pH 4 and enzyme added (enzyme/substrate ratio: 25/1,000, (w/w)). The solution was incubated at 44°C for 1.75 h. Ethanol was added to the hydrolysate to precipitate oligosaccharides with high degrees of polymerization overnight at 4°C. The suspension was then centrifuged and the supernatant was recovered, concentrated and then desalted using size exclusion chromatography on Sephadex G10 (77 cm ×44 mm). The column was equilibrated with degassed deionized water. RG-I oligosaccharides were loaded and eluted at a flow rate of 0.4 mL/min. Fractions collected were analyzed by high-performance anion-exchange chromatography (HPAEC). Appropriate fractions containing RG-I oligosaccharides were pooled and freeze-dried.

### Immunisation procedures and isolation of mAbs

The sugar beet RG-I oligosaccharides (10 mg) were coupled to BSA to produce the immunogen (Lees et al, 1996). The coupling efficiency was confirmed by phenol-sulfuric acid assay. Two male Wistar rats were injected with ∼200 μg of RG-I oligosaccharide-BSA in complete Freund’s adjuvant administered subcutaneously on day 0. The same amount was then administered with incomplete Freund’s adjuvant on days 21 and 62. On day 105 a rat was given a pre-fusion boost of the immunogen (1 mg in 1 mL PBS) injected intraperitoneally. The spleen were then harvested 3 days later and the lymphocytes were fused with myeloma cell line IR983F (Bazin, 1982). Hybridoma cells were selected by ELISA using the BSA-coupled oligosaccharides as the immobilised antigen.

### mAb characterization

LM6-M specificity was determined using ELISA (Cornuault et al., 2015) and epitope detection chromatography (EDC, Cornuault et al., 2014) on a range of plant materials: wheat arabinoxylan (Megazyme P-WAXYL), sugar beet de-branched arabinan (Megazyme P-DBAR), Tamarind xyloglucan (Megazyme P-XYGLN), sugar beet arabinan (Megazyme P-ARAB). Competitive-inhibition ELISAs of hapten recognition were performed using 1,5-arabinosyl oligosaccharides of DP2, 3, 5, 6, 7 and 9 (Megazyme O-ABI, O-ATI, O-APE, O AHE, O-AHP, O-ANO) as described in Marcus et al., 2008. In addition, susceptibility of the mAb to enzymatic action was investigated using α-L-arabinofuranosidase (*Aspergillus nidulans*, Megazyme E-ABFAN) and *endo*-1,4-β-galactanase (*Aspergillus niger*, Megazyme E-EGALN) on sugar beet arabinan (100μg/mL in PBS). For analysis of arabinan polysaccharides by epitope detection chromatography (EDC) Arabidopsis plants were grown at 20° C under 16 h light/8 h dark cycle for 30 days. Inflorescence stems were harvested and frozen in liquid nitrogen before being freeze-dried and homogenized. Alcohol insoluble residues were prepared by treating the sample with increasing amounts of ethanol (70, 80, 90 and 100%), followed by 100% acetone and methanol:choloroform (2:3) for 1 h in each solution. The remaining material was dried and the pectic cell wall polymers were extracted using 50 mM CDTA as detailed (Cornuault et al., 2014). Indirect immunofluorescence labelling of plant materials was carried out as described (Cornuault et al., 2015).

## Results & Discussion

In order to extend the repertoire of available mAbs to RG-I polysaccharides a series of RG-I-enriched preparations were isolated from potato tubers (Buffetto et al. 2015; Cornuault et al. 2015) and, as reported here, from sugar beet roots to prepare immunogens. The impetus of the isolation procedure for sugar beet RG-I oligosaccharides was to obtain mAbs to epitopes of the RG-I backbone/base of the arabinan side chains. Sugar beet pulp is an established source of arabinan-rich RG-I polysaccharides (Oosterveld et al., 2000). Extracted polysaccharides were separated by anion-exchange chromatography (Fig. 1a) and pools 1 and 2, enriched in neutral sugars reflecting high levels of RG-I (Table 1) were combined in an RG-I pool which was then hydrolysed with HCl to reduce the length of side chains.

**Table 1.** Sugar composition of the pectin, purified RG-I and RG-I oligosaccharides from sugar beet in mol%.

|  | GalA | Rha | Gal | Ara |
| --- | --- | --- | --- | --- |
| **Pectin extracted** | 36.00 | 7.00 | 10.00 | 47.00 |
| **RG-I (1 & 2)** | 14.00 | 9.00 | 13.00 | 64.00 |
| **RG-I oligosaccharides** | 23.00 | 19.00 | 29.00 | 29.00 |

**Figure 1.**
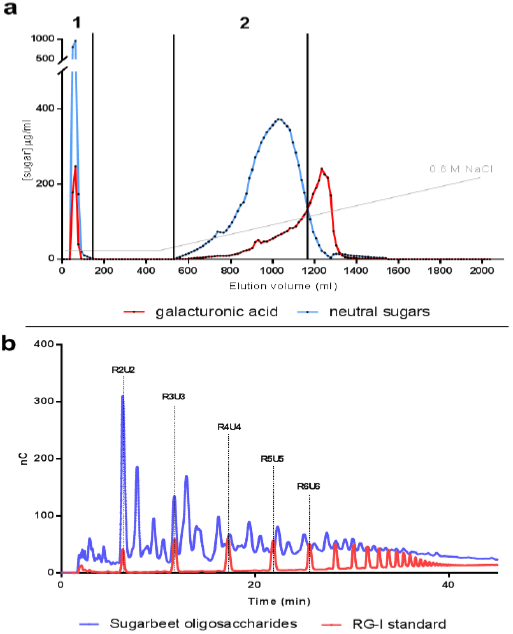
Preparation of sugar beet RG-I oligosaccharides.(**a)**Anion-exchange chromatography of isolated sugar beet pectin, composition of acidic and neutral sugars determined by automated *m*-hydroxybiphenyl and orcinol methods respectively. (**b)** High performance chromatography profile of the obtained sugar beet oligosaccharides compared to RG-I backbone oligosaccharides standards.

The isolated RG-I preparation was then partially digested by rhamnogalacturonase to obtain a range of oligosaccharides (Fig. 1b). These RG-I oligosaccharides were used to prepare an immunogen by coupling to BSA prior to standard rat immunisation and rat hybridoma cell line isolation protocols. However, out of the twenty antibody-secreting cell lines obtained subsequent to immunisation with RG-I-BSA none were found to bind to the decorated RG-I backbone, after ELISA screening on a range of RG-I preparations, but instead several bound to linear 1,5-L-arabinan. This raises questions of the differences in immunogenicity between ramified or linear structures, acidic and neutral carbohydrate structures and immunodominance that are not addressed further here. It is also possible that structural features of the RG-I backbone and aspects of steric hindrance within branched polymers influence both the generation of antibodies within immune systems and/or the selection of antibodies during hybridoma screening.

One mAb was found to bind to 1,5-L-arabinan in ELISA screens with higher signals than the widely used arabinan mAb LM6 (Willats et al., 1998) and was characterized further, alongside LM6, and was found to have a closely related specificity. This mAb has an IgM isotype and as LM6 is an IgG the new arabinan-directed mAb was designated LM6-M. Figure 2a shows the comparative binding of LM6 and LM6-M hybridoma supernatants to four polysaccharides and indicates the strong signal obtained from the arabinan polysaccharides with LM6-M and no recognition of arabinoxylan or xyloglucan. Binding of LM6-M and LM6 to sugar beet arabinan were equally abrogated by α-L-arabinofuranosidase (Fig. 2b). Hapten-inhibition ELISA studies with arabino-oligosaccharides indicated that effective recognition by LM6-M of 1,5-arabinosyl residues is obtained with three residues, as for LM6 (Fig. 3) although the IC50 for the arabinotriose is ∼7-fold higher for LM6-M than LM6.

**Figure 3.**
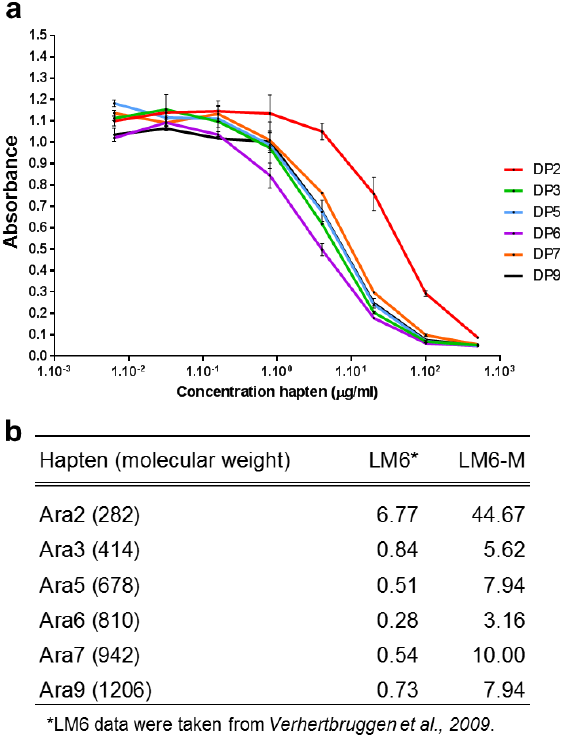
ELISA hapten inhibition analysis of LM6-M using a range of arabinan oligosaccharides ranging in degree of polymerisation from 2 to 9. (**a)** Capacity of arabino-1,5-saccharide haptens to inhibit the binding of LM6-M. (**b)** Table showing the IC50 (50% binding inhibition) values for both LM6 and LM6-M recorded in ¼;g/ml against the different arabinan oligosaccharides. Values for LM6 taken from Verhertbruggen et al., 2009.

**Figure 2.**
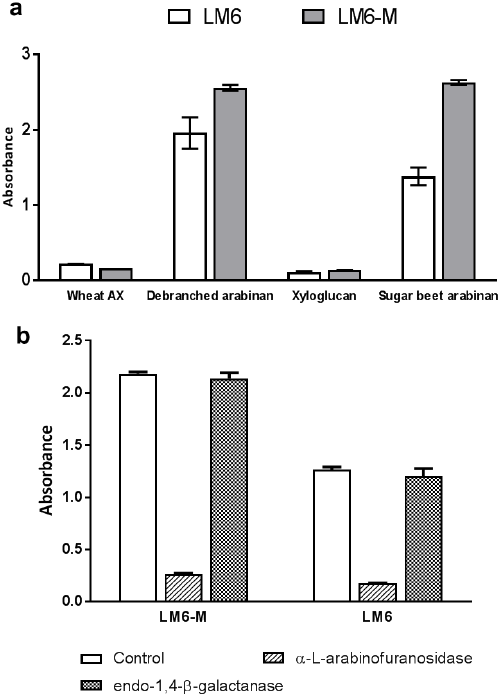
**(a)** ELISA of LM6 and LM6-M mAbs binding to wheat arabinoxylan (AX), debranched arabinan, xyloglucan and sugar beet arabinan. Polysaccharides all coated on to plates at 50 μg/ml.(**b**) Effect of α-L-arabinofuranosidase and endo-1,4-β-galactanase enzyme (both at 100 μg/ml) digestion on the binding of LM6 and LM6-M to sugar beet arabinan. Error bars = SE, n = 3.

Use of LM6-M in conjunction with anion-exchange-epitope detection chromatography (AE-EDC, Cornuault et al., 2014) confirmed the increased detection of arabinan by this mAb relative to LM6 (Fig 4) for both sugar beet arabinan and also a CDTA-extract of Arabidopsis inflorescence stem cell walls. Both profiles indicated recognition of two peaks by LM6-M and LM6. In the case of the Arabidopsis stem extract the major peak coincided with pectic HG (Cornuault et al., 2014) and LM6-M and LM6 additionally detected an early eluting, less acidic, shoulder that was not detected by LM13 – a mAb requiring longer arabinan chains for recognition (Verhertbruggen et al., 2009). A similar pattern is observed for the sugar beet arabinan where two distinct peaks are recognized by the three mAbs with LM13 binding preferentially to the second, later eluting, one. It is interesting to note that the AE-EDC profile for sugar beet arabinan is different to that obtained from Arabidopsis stem with the arabinan appearing earlier in the elution gradient. There is no indication of neutral molecules in the isolated arabinan preparation indicating that the polysaccharides are still attached to acidic pectin, most probably RG-I backbone.

**Figure 4.**
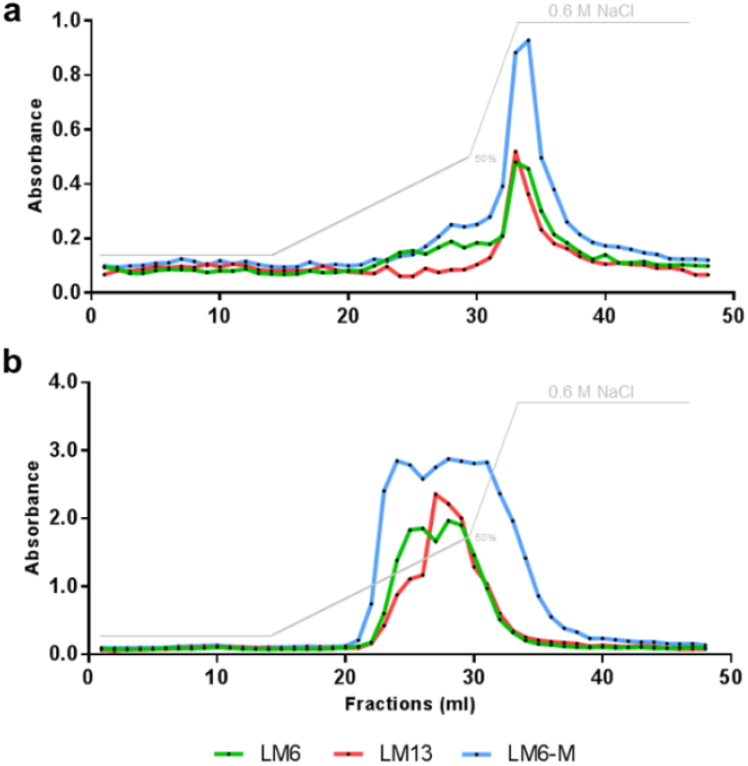
AE-EDC profiles of arabinan epitopes in Arabidopsis stem CDTA extract and sugar beet arabinan using 1,5-arabinan probes LM6-M, LM6 and LM13. Salt elution gradient shown by faint grey line.

Iindirect immunofluorescence procedures with LM6-M indicated that it bound effectively to plant materials as shown for transverse sections of Arabidopsis inflorescence stem (Fig. 5). In this case the LM6-M epitope was particularly abundant in epidermal cell walls which has previously been shown for pectic arabinan (Verhertbruggen et al., 2009).

**Figure 5.**
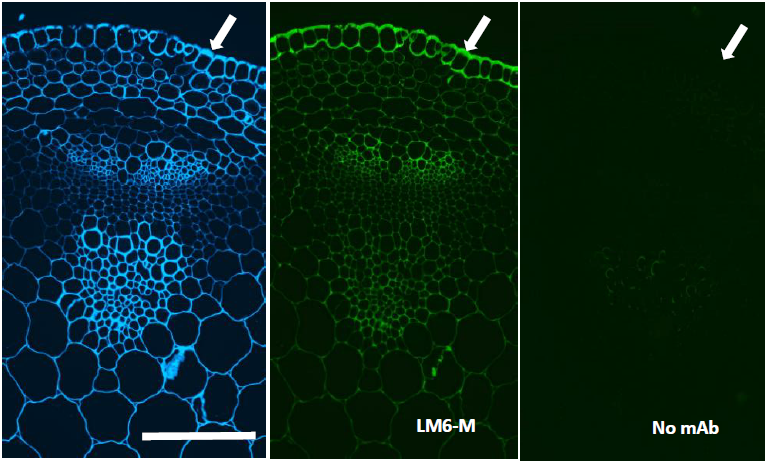
Indirect immunofluorescence detection of the LM6-M epitope in a transverse section of the Arabidopsis inflorescence stem. LM6-M binds to all cell walls and strongly to epidermal cell walls. Blue indicates Calcofluor White staining of the same section as shown for LM6-M fluorescence and an equivalent section shows the no mAb control. Arrows indicate the stem surface and outer epidermal cell wall. Scale bar = 10 μm.

## Conclusion

LM6-M is a high avidity rat mAb directed to 1,5-arabinosyl residues and its high detection capability makes is useful for the analysis of this structural feature of RG-I in a range of antibody-based techniques. It is probable that the observed signal enhancement by LM6-M over LM6 is the result of both increased multivalent capacity of IgM immunoglobulins over IgG immunoglobulins and also a signal amplification obtained by increased binding of secondary antibodies to larger IgM molecules.

## Acknowledgements

We thank Alex Keen for help with the characterization of LM6-M. This work was supported by the European Union Seventh Framework Programme (FP7 2007-2013) under the WallTraC project (Grant Agreement number 263916). (This article reflects the authors’ views only and the European Union is not liable for any use that may be made of the information contained herein). The work was also supported by the United Kingdom Biotechnology and Biological Research Council (BBSRC, grant BB/K017489/1).

